# BrainRBPedia: a resource for RNA-binding proteins relevant to neurodevelopmental disorders

**DOI:** 10.1101/2023.06.07.542483

**Authors:** Kara Han, Michael Wainberg, John A. Calarco, Craig A. Smibert, Howard D. Lipshitz, Hyun O. Lee, Shreejoy J Tripathy

## Abstract

RNA-binding proteins (RBPs) are crucial players in the post-transcriptional regulation of mRNA and play major roles in ensuring proper neuronal development and function. Deficits in RBP function have been implicated in a number of neurodevelopmental disorders including autism spectrum disorder (ASD) and intellectual disability (ID), yet we lack resources that integrate current knowledge of RBP function, tissue expression, and disease association in one place to aid in their experimental characterization. Here we introduce BrainRBPedia – a database of 1072 RBPs with both disease annotations for neurodevelopmental disorders and functional annotations relevant to these disorders, including loss-of-function intolerance and expression specificity to the brain, neurons, and neuronal development. Using these functional annotations, we develop a machine learning model to prioritize RBPs likely to be involved in ASD and ID. Our model indicates that RBPs with high loss-of-function intolerance and those upregulated during neuronal differentiation are disproportionately likely to contribute to ASD and ID etiology. In summary, BrainRBPedia comprises a unique resource for researchers interested in the experimental characterization of RBPs in relation to neurodevelopmental disorders and suggests functional signatures of RBPs likely to play a role in neurodevelopment.

## INTRODUCTION

RNA-binding proteins (RBPs) bind RNA via one or more RNA-binding domains (RBDs) and play crucial roles in every step of post-transcriptional control of gene expression (Hentze et al., 2018). These include a diverse array of processes such as pre-mRNA splicing, editing, polyadenylation, stability, transport, localization, and translation. These processes are especially important in the intricate networks of gene expression control during neurodevelopment, and numerous studies have now identified key RBPs involved in each of these steps.

As examples, murine PTBP1 acts as a regulatory switch of alternative splicing programs during neuronal cell differentiation (Boutz et al., 2007) and its human ortholog PTBP2 governs axonogenesis-associated splicing and controls the fate of early axon formation (M. Zhang et al., 2019). RNA editing by adenosine deaminase acting on RNAs (ADARs) is particularly active in the brain (Mehler & Mattick, 2007), and this activity is proposed to contribute to the complexity of the nervous system. RBPs also control alternative polyadenylation. For example, CPEB1 has been implicated in synaptic plasticity (Alarcon et al., 2004) and dendritic arborization (Bestman & Cline, 2008), and promotes 3’-UTR shortening events (Bava et al., 2013). The RBP, fragile X mental retardation protein (FMRP) functions in the transport of RNAs to the synapse and modulates the translation of numerous synaptic proteins (Antar et al., 2005).

Dysregulation of RBPs during neurodevelopment also affects proper neuronal function and underlies a broad spectrum of neuropsychiatric disorders, particularly autism spectrum disorders (ASD) and intellectual disability (ID). For example, deficits in FMRP function lead to fragile X syndrome (FXS), the leading monogenic cause of ASD and inherited ID (Gantois et al., 2017). The TNRC6B RBP functions in microRNA-mediated gene silencing; pathogenic variants cause a genetic disorder characterized by ID and other autism-related features (Granadillo et al., 2020). The DDX3X RNA helicase plays a critical role in neurite outgrowth, synaptogenesis, and proliferation and differentiation of cortical neural progenitors in the brain (Lennox et al., 2020); mutations in *DDX3X* cause one of the most common genetic ID syndromes in females (Snijders Blok et al., 2015).

Despite the importance of RBPs to neurodevelopmental disorders, we lack resources that bring together existing knowledge on RBPs that are highly expressed in neuronal cell types, their association with disease, and potential functions. While existing RBP databases focus on generic characteristics like their motifs and binding affinities (Caudron-Herger et al., 2021; Cook et al., 2011; Liao et al., 2020), no databases exist, to our knowledge, that enable researchers interested in experimentally characterizing the roles of RBPs in neurodevelopmental disorders to help prioritize their efforts. Here, we introduce BrainRBPedia — a database of RBPs annotated for functions relevant to neurodevelopmental disorders, including protein domains, brain and neuron specificity, mutational constraint, and neurodevelopmental disorder susceptibility. BrainRBPedia annotations were derived by leveraging and re-analyzing public ‘omics datasets and gene-disease annotations. In addition, we developed a machine learning model using these annotations to predict and prioritize ASD and ID candidate RBPs. This model also identifies functional annotations in BrainRBPedia that are predictive of whether an RBP is known to be associated with ASD and ID. BrainRBPedia comprises a unique resource for prioritizing neurodevelopmental disorder-related RBPs for experimental follow-up.

## MATERIALS AND METHODS

### A Curated list of RNA Binding Proteins

As the basis of BrainRBPedia, we used a curated list of 1072 RBPs and their associated Ensembl IDs and gene symbols (Sundararaman et al., 2016). These proteins were annotated as RBPs based on the union of two gene lists: genes with canonical RNA binding domains (N = 476) (Castello et al., 2012) and genes identified experimentally as RBP candidates via interactome capture in HeLa cells (N = 845) (Gerstberger et al., 2014).

Corresponding mouse gene symbols were acquired using the human2mouse function provided in the homologene R package (Mancarci, 2019).

### Protein domains and canonical RBDs

BrainRBPedia’s “Protein domains” column lists each RBP’s protein domains. Protein domains were obtained from the “interpro” attribute provided in the Ensembl BioMart database via the biomaRt R package (Durinck et al., 2009). Protein domains are only listed once even if they are present multiple times within an RBP.

BrainRBPedia’s “Contains canonical RBDs” column denotes which RBPs contain one or more canonical RNA binding domains (RBDs). An RBP was deemed to have a canonical RBD if any of its protein domains listed in BioMart had the same Pfam and InterPro ID as one of the RBDs listed in CIS-BP-RNA (Ray et al., 2013) or one of our manually curated RBDs (see Table S4).

### Cell and tissue expression specificity

#### Neuron Enrichment

BrainRBPedia’s “Neuron enrichment (mouse)” column summarizes each RBP’s gene expression enrichment in neurons relative to non-neurons in mouse models. To create this column, we used a mouse cortical neuron single-cell RNA-seq dataset generated by the Allen Institute for Brain Science (Tasic et al., 2018), denoted hereafter as the Tasic 2018 dataset. Tasic 2018 comprises 15,413 cells from the primary visual cortex (VISp) and 10,068 cells from the anterior lateral motor cortex (ALM). A mouse dataset was chosen here to acquire mRNA expression from the entire cell as opposed to just the nucleus.

We used the Seurat R package (Hao et al., 2021) workflow to process the data and perform the neuronal differential expression tests on the Tasic 2018 dataset. Specifically, we applied the Seurat FindMarkers function with default settings to find genes distinguishing neurons (GABAergic or Glutamatergic) from non-neurons (all other cell classes), yielding the average log_2_ fold change between the neuronal and non-neuronal groups. Positive values indicate that the gene is more highly expressed in neurons than in non-neurons. For the few human RBPs with multiple mouse orthologs, we kept the mouse ortholog with the highest mouse neuron enrichment value.

Similarly, BrainRBPedia includes a “Neuron enrichment (human)” column, which summarizes RBPs’ gene expression enrichment in human neurons relative to non-neurons. We used single-nucleus RNA-seq obtained from the middle temporal gyrus of the human cortex (Hodge et al., 2019). The construction of this column followed the same workflow as the “Neuron enrichment (mouse)” column using the Seurat R package.

#### Brain Enrichment

BrainRBPedia includes a “Brain enrichment” column, which summarizes each RBP’s gene expression enrichment in the brain relative to other tissues in the Genotype-Tissue Expression (GTEx) dataset (GTEx Consortium, 2013). Specifically, we used the *Median gene-level Transcripts Per Million (TPM) by tissue* RNA-seq data found in GTEx Analysis V8 to calculate each RBP’s log_2_ fold difference in TPM between brain and non-brain tissues. Similar to Neuron enrichment, positive values indicate that the RBP is more highly expressed in the brain than in non-brain tissues.

#### Neuron Developmental Enrichment

To study RBP gene expression during neuronal differentiation and polarization, we made use of the transcriptomic profiles of human induced pluripotent stem cell (hiPSC)-derived neurons sampled at early stages of neuron development, specifically, days 1, 3, and 7 post-differentiation (Lindhout et al., 2020). These sampling time points correspond to three well-defined stages of neuronal differentiation and polarization: the formation of small processes (stage 1, day 1), multiple neurites (stage 2, day 3), and cell polarization, wherein one neurite is specified as the axon (stage 3, day 7). We specifically made use of the *day3 vs. day1* and *day7 vs. day1* differential gene expression information from Lindhout et al. as the “Neuron developmental enrichment (early)” and “Neuron developmental enrichment (late)” columns in BrainRBPedia.

#### Protein Expression (Caudate/Cerebral Cortex/Hippocampus)

BrainRBPedia includes three “Protein expression” columns with each RBP’s protein expression in three brain regions: caudate nucleus, cerebral cortex and hippocampus. Protein expression was measured using immunohistochemistry and tissue microarrays and obtained from The Tissue Atlas, part of The Human Protein Atlas project (Uhlén et al., 2015). For each brain region, an RBP’s protein expression was assigned by the original authors one of four possible values: High, Medium, Low, and Not detected.

### Mutational constraint

BrainRBPedia includes the metric of mutational constraint defined as the “Probability of loss-of-function intolerance (pLI)” column obtained from the *pLoF Metrics by Gene TSV* found in the Genome Aggregation Database (gnomAD) v2.1.1 dataset, which contains data from 125,748 exomes and 15,708 whole genomes mapped to the GRCh37/hg19 reference sequence (Karczewski et al., 2020). The pLI score is a widely used metric of intolerance to loss-of-function mutations and ranges from 0 (no evidence of loss-of-function intolerance) to 1 (strong evidence of loss-of-function intolerance). For downstream analyses making use of these pLI scores, we chose to group scores into four ranges as we have performed previously (Wainberg et al., 2022): low (<0.50), medium (0.50 to 0.90), high (0.90 to 0.99), and extreme (0.99 to 1).

### Neurodevelopmental disorder genes

The annotation of genes implicated in ASD susceptibility was downloaded from the Human Gene Module of the Simons Foundation Autism Research Initiative (SFARI) Gene database version 2018 Q4 (Abrahams et al., 2013).

The “Intellectual disability susceptibility” column was obtained from the “Intellectual disability” (version 4.12) panel in Genomic England’s PanelApp (Martin et al., 2019), where gene-disease annotations are reviewed by experts in the worldwide scientific community. We only selected genes with “Green” (high level of evidence for gene-disease association) and “Amber” (moderate level of evidence) internal ratings to ensure the quality of gene-disease annotations.

### Predictive model construction for identifying informative features and novel ASD and ID RBPs

We used penalized multivariate logistic regression (LR) to predict whether or not each RBP was present in the SFARI ASD database or the Genomics England “Intellectual disability” gene panel based on its functional annotations in BrainRBPedia, as described above: Contains canonical RBDs, Neuron enrichment (human), Neuron enrichment (mouse), Brain enrichment, Neuron development enrichment (early), Neuron development enrichment (late), and pLI.

To account for the fact that genes with pLI scores very close to 1 are disproportionately relevant to neuropsychiatric disorders (Wainberg et al., 2022), the pLI score was modelled as three binary variables – “Medium pLI” (whether or not an RBP’s pLI score is between 0.5 and 0.9), “High pLI” (between 0.9 and 0.99), and “Extreme pLI” (between 0.99 and 1).

Missing values in the continuous functional annotations were replaced with the mean of the non-missing values for that variable. To select relevant functional annotations and classify candidate ASD/ID genes, we fit a multivariate least absolute shrinkage and selection operator (LASSO) logistic regression model on the 1072 RBPs using the *cv.glmnet* function in the *glmnet* R package (Friedman et al., 2010). LASSO reduces model complexity by shrinking the beta coefficients of unimportant variables and prevents overfitting that may result from simple LR models.

We used nested cross-validation, with an outer five-fold cross-validation to evaluate predictive performance and an inner five-fold cross-validation within each of the outer folds to select the L1 penalty. To account for class imbalance (i.e., most RBPs not having a known link to ASD/ID), we ensured that each fold contained the same ratio of positive to negative examples as the full dataset, and also unweighted positive examples in direct proportion to their scarcity using the *weights* argument to *cv.glmnet*. We used *lambda.1se* obtained from the cross-validation fit to extract the beta coefficients for each of the five folds. *lambda.1se* denotes the largest L1 penalty for which the validation error was within 1 standard error of the minimum error across all L1 penalties.

We evaluated model performance by calculating the area under the receiver operating characteristic (ROC) curve (AUC) and the area under the precision-recall curve (AUPRC) using the *ROCR* R package (Sing et al., 2005).

## RESULTS

### A database of brain-related functional annotations for RNA Binding Proteins

To examine the relationship between RBPs and neuronal development and disorders, we annotated 1072 human RNA-binding proteins (RBPs) with multiple functional annotations from publicly available data sources from mice and humans (Figure 1A). These annotations encompass whether an RBP contains a canonical RNA-binding domain (RBD), shows enriched expression in brain tissues relative to non-brain tissues, in neurons relative to non-neurons, or in specific phases of neuronal development, whether the RBP’s coding sequence is depleted for loss-of-function mutations in the general population (loss-of-function intolerant), or is associated with neurodevelopmental disorders (Figure 1). The complete results of these analyses are packaged into a database called BrainRBPedia in Table S2.

**Figure 1:**
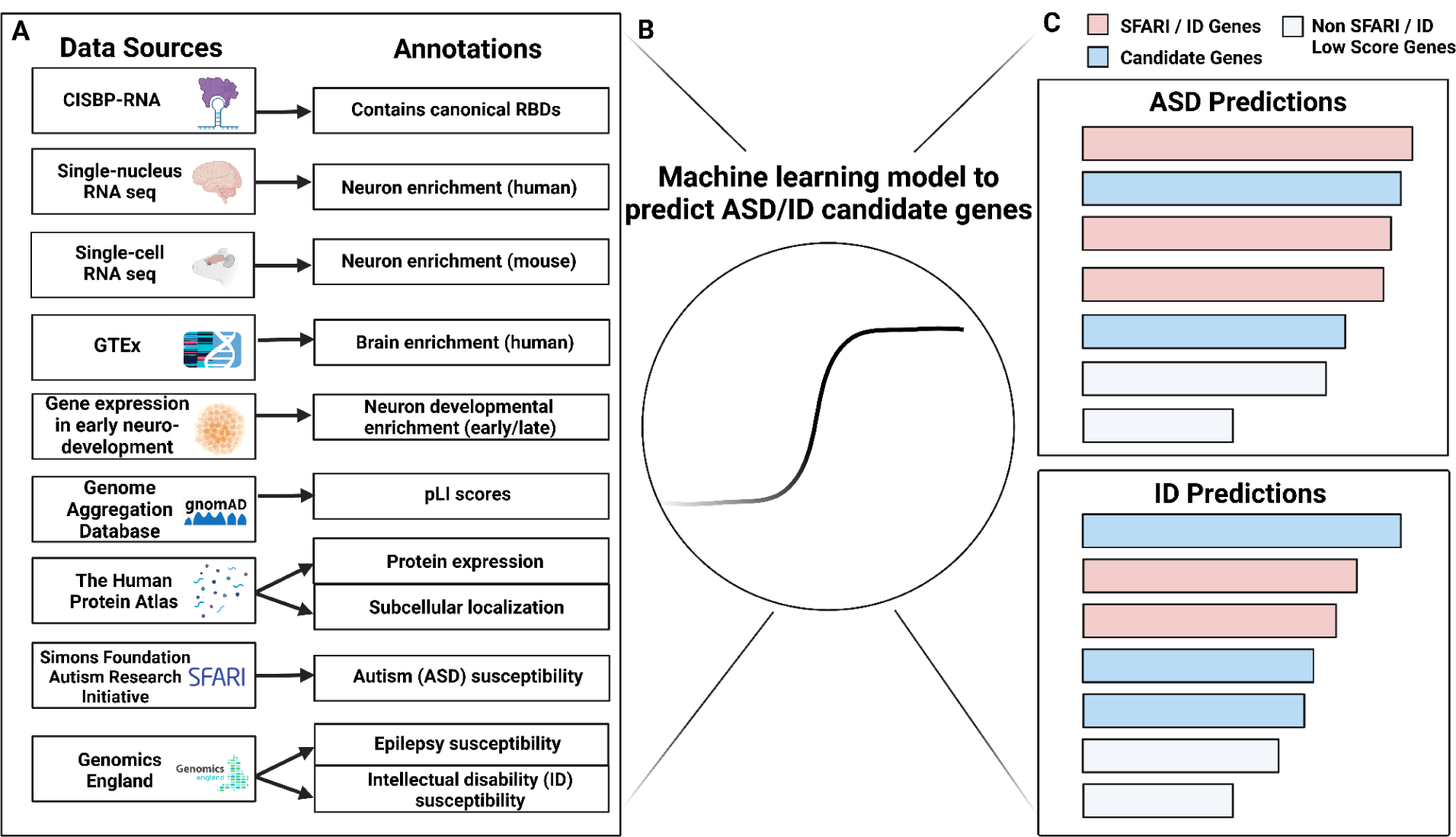
An overview of BrainRBPedia. (A) Multiple publicly accessible data sources were used to generate the functional annotations in BrainRBPedia for 1072 human RBPs. (B) Selected functional annotations were used to construct a machine learning model that predicts ASD and ID candidate genes. (C) ASD and ID candidate gene predictions from the machine learning model. The length of each box indicates the likelihood of ASD susceptibility.

Distributions of key functional annotations from BrainRBPedia are provided in Figure 2. Figure 2A shows the enrichment of gene expression of RBPs and other protein-coding genes in human adult neurons relative to non-neurons, using single-nucleus RNAseq data from the neocortex (Hodge et al., 2019). We found that though non-RBP protein-coding genes are generally disenriched in neurons relative to non-neurons, genes encoding RBPs show significantly reduced enrichment in neurons compared to non-RBP coding genes (asymptotic two-sample KS test: D = 0.16, p-value < 2.2e-16). The bimodal nature of human neuron enrichment observed in both RBPs and other protein-coding genes suggests that both RBPs and non-RBPs have a set of genes that show enriched in neurons and a set that are not. We next examined whether this trend is consistent in the brain in comparison to other non-brain tissues. We quantified the gene expression of RBPs and other protein-coding genes in brain tissues relative to non-brain tissues using RNAseq data from GTEx (GTEx Consortium, 2013) in Figure 2B. It can be seen that RBPs are less enriched in the brain relative to non-brain tissues than non-RBP coding genes (KS test: D = 0.55, p-value < 2.2e-16), similar to the trend observed with neurons vs. non-neurons. These findings indicate that RBPs are generally disenriched in adult neurons compared to non-neurons and brain tissues compared to non-brain tissues compared to other protein-coding genes.

**Figure 2:**
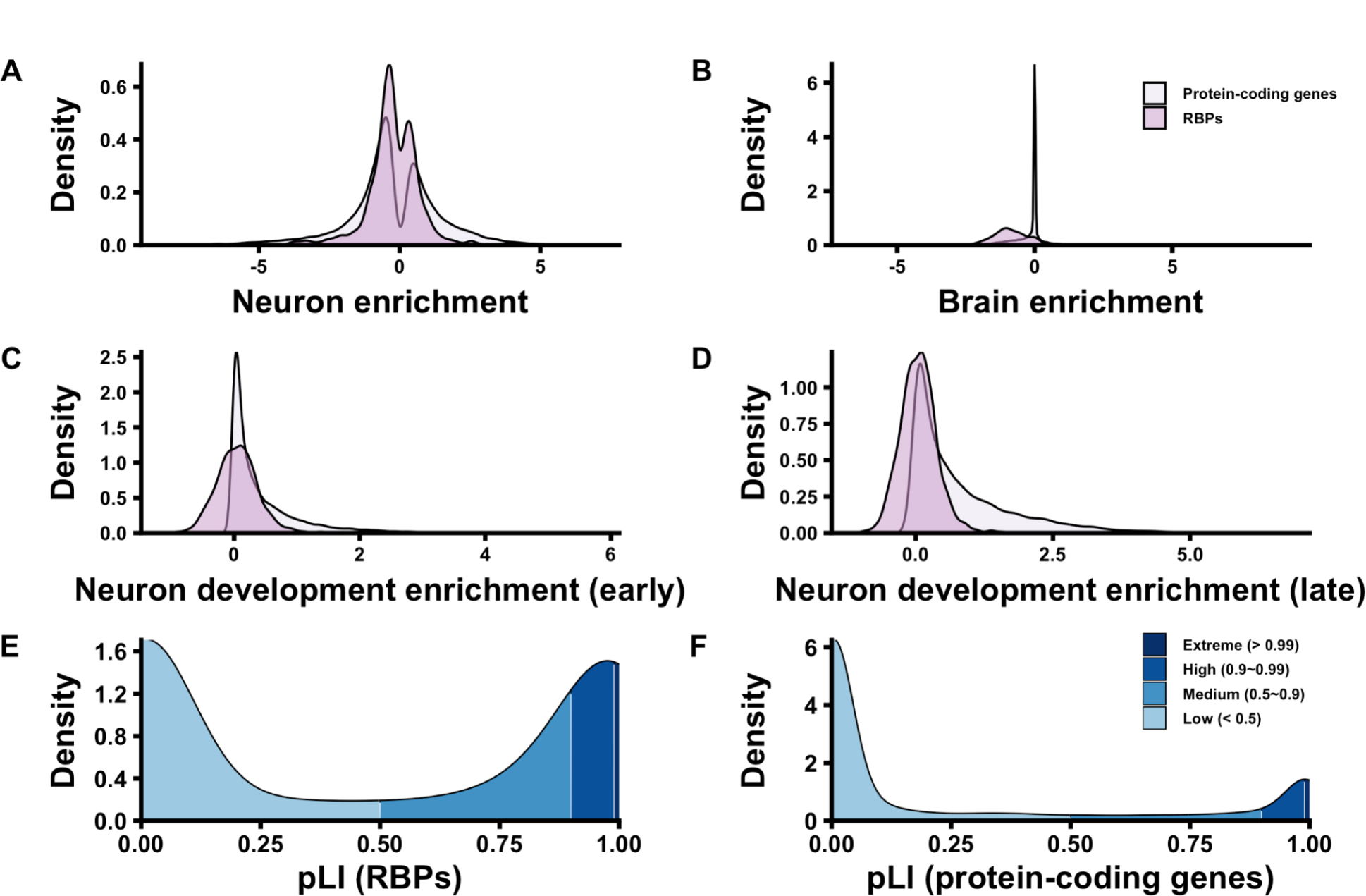
Distribution of key BrainRBPedia functional annotations. (A-D) Density plots of neuron enrichment values, relative to non-neurons, in human adult neurons (A), in adult brain tissues relative to non-brain tissues (B), or early and late neuron developmental enrichment values, based on human hiPSC-derived neuron cultures (C-D) for RBPs and non-RBP coding genes. The early enrichment is measured by the day 3 vs. day 1 differential expression, while the late enrichment is computed using the day 7 vs. day 1 differential expression change. (E-F) Density plots of pLI scores for RBPs and non-RBP coding genes, respectively, with four ranges: low (<0.50), medium (0.50 to 0.90), high (0.90 to 0.99), and extreme (0.99 to 1).

To understand how gene expression profiles change over the course of neuronal development, we examined the relative enrichment of RBPs and non-RBP coding genes using transcriptomic profiles from human induced pluripotent stem cells (hiPSC) sampled at different stages of neuronal differentiation (Figure 2C-D). Globally, RBPs show more enriched expression at an earlier stage of neuronal development, with 548 genes upregulated at day 3 of differentiation relative to day 1 and 433 genes downregulated at day 3 relative to day 1. The expression of RBPs becomes less enriched at day 7 compared to day 1, where 309 genes are upregulated and 672 genes are downregulated. Compared to non-RBP coding genes, RBPs showed less enriched expression during both early- and late-stages of neuron development (KS test early stage: D = 0.44, p-value < 2.2e-16; KS test late stage: D = 0.69, p-value < 2.2e-16).

Next, we analyzed if RBPs as a group are under strong mutational constraint by assessing their loss-of-function as computed from large human exome- and genome-wide sequencing datasets from control individuals (Karczewski et al., 2020). We specifically used pLI scores (probability of loss-of-function intolerance) and grouped scores into four ranges: low (<0.50), medium (0.50 to 0.90), high (0.90 to 0.99), and extreme (0.99 to 1). We found that most RBPs have either very low pLI scores, with 50.7% of RBPs demonstrating low mutational constraint (pLI < 0.5), or very high or extreme pLI scores (pLI > 0.99), indicating very strong mutational constraint (26.0% of RBPs). In contrast, only 10.3% of non-RBP coding genes (see Figure 2F) showed very strong mutational constraints (pLI > 0.99), indicating that RBPs are considerably more likely to be mutationally constrained and be loss-of-function intolerant.

In summary, compared to non-RBP coding genes, RBPs showed less enrichment in neurons relative to non-neurons, in brain relative to non-brain tissues, and in both early and late neuron developmental stages. However, RBPs are under very strong mutational constraint in comparison to other protein-coding genes in the genome.

### A predictive model of neurodevelopmental-disorder relevant RBPs

Building on these analyses of RBPs in neurodevelopment, we next aimed to use a machine-learning framework to predict RBPs relevant to autism spectrum disorders (ASD) and intellectual disability (ID) using the annotations for each RBP in BrainRBPedia. In addition to identifying potentially novel RBPs related to these disorders, we also aimed to understand which underlying RBP annotations (e.g., loss of function intolerance, gene expression enrichment in neurodevelopment, etc.) were most predictive of a gene being associated with each of these diseases. We made use of the disease specific annotations from SFARI for ASD-related genes (Abrahams et al., 2013) and Genomics England for ID-related genes (Martin et al., 2019). In total, there were 81 ASD-related RBP genes (991 non-ASD RBPs) and 148 ID-related RBP genes (924 non-ID RBP genes). We used penalized multivariate logistic regression (Figure 1B) and validated the model’s performance using five-fold cross-validation.

The selected set of features in each of the five model runs and their odds ratios are presented in Figure 3. In the model predicting ASD-related RBP genes, the most informative features were loss-of-function intolerance, and in particular, whether RBP genes had pLI scores in the extreme (0.99 to 1) or high categories (0.90 to 0.99), and also the degree to which an RBP gene was upregulated over later stages of neuron development (i.e., upregulated between day 7 versus day 1, days in vitro). After adjusting for the presence of other predictors used in the model, RBPs with extreme and high pLI scores (i.e., pLI between 0.90 and 1) are, on average, 4.00 and 1.77 times more likely to be associated with ASD than those without extreme and high pLI scores. We also found that RBPs that are more highly expressed at day 7 relative to day 1 during neuronal development are associated with a 23% increase in the odds of ASD susceptibility. Intriguingly, no other RBP features (e.g., whether an RBP contains canonical RNA binding domains or whether an RBP is enriched in the brain relative to other tissues) were consistently predictive of ASD gene susceptibility. Our findings suggest that loss-of-function intolerance and differential gene expression during neuron development are statistically predictive factors for ASD gene etiology.

**Figure 3:**
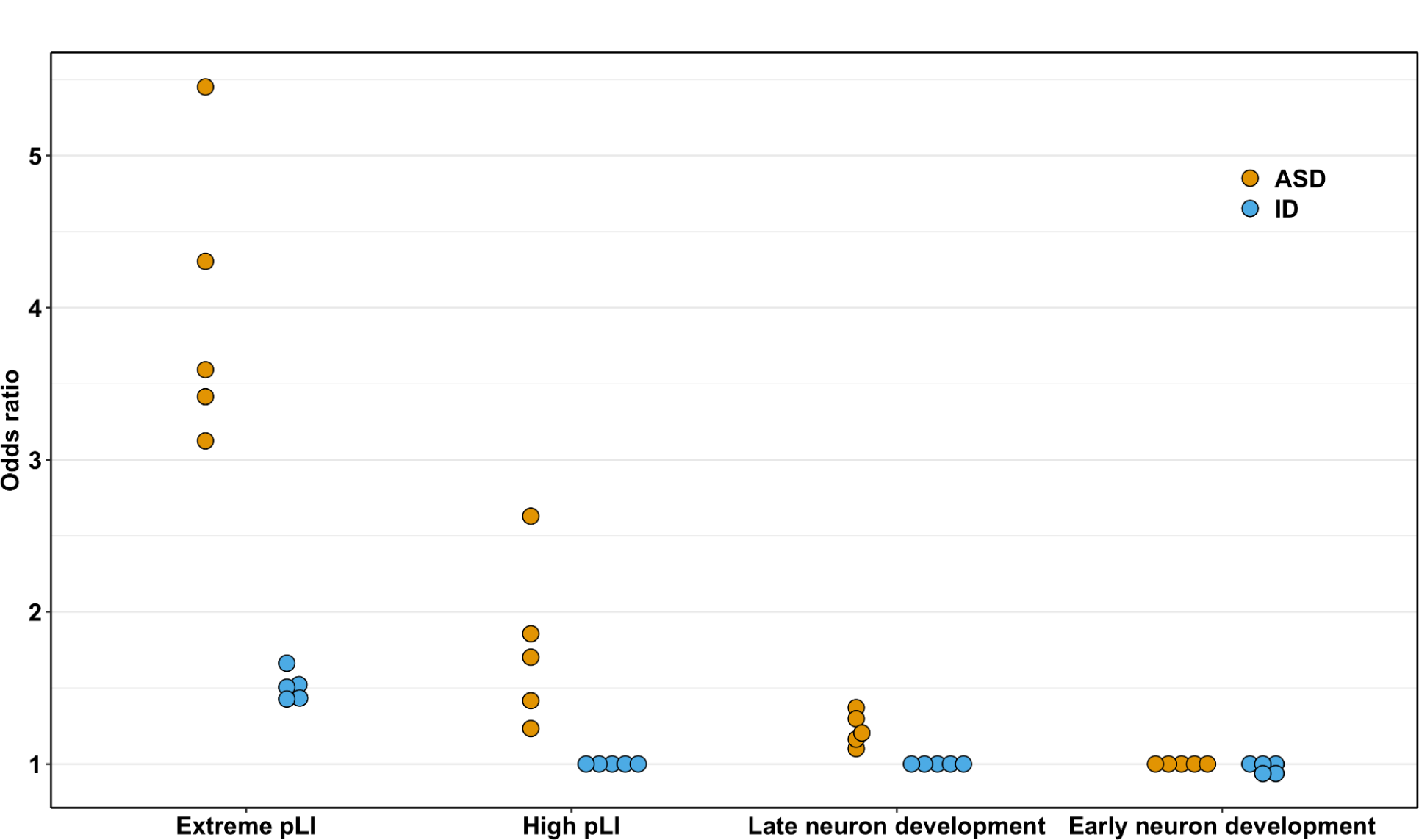
Informative RBP features for ASD and ID susceptibility gene prediction obtained from the penalized multivariate logistic regression models. For each predictor variable (x-axis), the five odds ratios obtained from the five-fold cross validations are shown for the ASD (blue) and ID (yellow) models. Odds ratios indicate the relative importance of each feature where an odds ratio of 1 indicates the feature is not predictive for a particular cross validation fold. Features never chosen by either model (e.g., whether an RBP has canonical RNA-binding domains) are not shown.

In ID-susceptibility gene prediction, the features of Extreme pLI and Neuron developmental enrichment early (i.e., upregulated between day 3 versus day 1 days in vitro) were statistically predictive, as indicated by their selection across different model runs. Similar to ASD susceptibility prediction, RBPs with extreme pLI scores (i.e., pLI between 0.99 and 1) were, on average, 1.51 times more likely to be implicated in ID than those without extreme pLI scores. On the other hand, each additional one-unit increase of an RBP’s differential expression at day 3 compared to day 1 during neuronal development is associated with a 3% decrease in the odds of its ID susceptibility.

In general, RBPs with extreme pLI scores are most likely to be associated with ASD and ID among all predictors in both models, suggesting that the intolerance of an RBP to loss-of-function mutations is strongly predictive of its relevance to ASD and ID etiology. Additionally, the upregulation of RBPs during specific critical stages of neuronal development, including the formation of small processes (day 3) and multiple neurites (day 7), appears to play a role in their association with ID and ASD, respectively, albeit to a lesser extent than that seen with extreme or high pLI scores.

To quantify the prediction accuracy of each model, we show five-fold model prediction accuracies for each of the ASD- and ID-RBP gene prediction tasks, shown as receiver-operating characteristic (ROC) and precision-recall (PR) curves in Figure 4. The ASD model (Figure 4A) showed a comparatively good prediction performance, with area under the curve values (AUCs) ranging from 0.72 to 0.89 across each of the five cross-validation folds (with differences between folds due to randomness in how RBP genes are selected for test and training sets), indicating that a randomly selected RBP in SFARI has a 72-89% chance of receiving a higher prediction score from the model than a randomly selected non-SFARI RBP. Similarly, AUPRCs for the ASD model ranged from 0.22 to 0.41 (Figure 4C), compared to a baseline of 0.076 (the fraction of the RBPs that are in SFARI). ROC AUCs for the ID model suggest that a randomly selected ID-susceptible RBP has a 58-66% chance of being ranked higher than non-ID RBPs (Figure 4B). AUPRCs ranged from 0.15 to 0.24, compared to a baseline of 0.138 (the fraction of the RBPs with ID associations). In summary, these approaches demonstrate that ASD- and ID-related RBPs can be predicted by machine learning algorithms using simple features.

**Figure 4:**
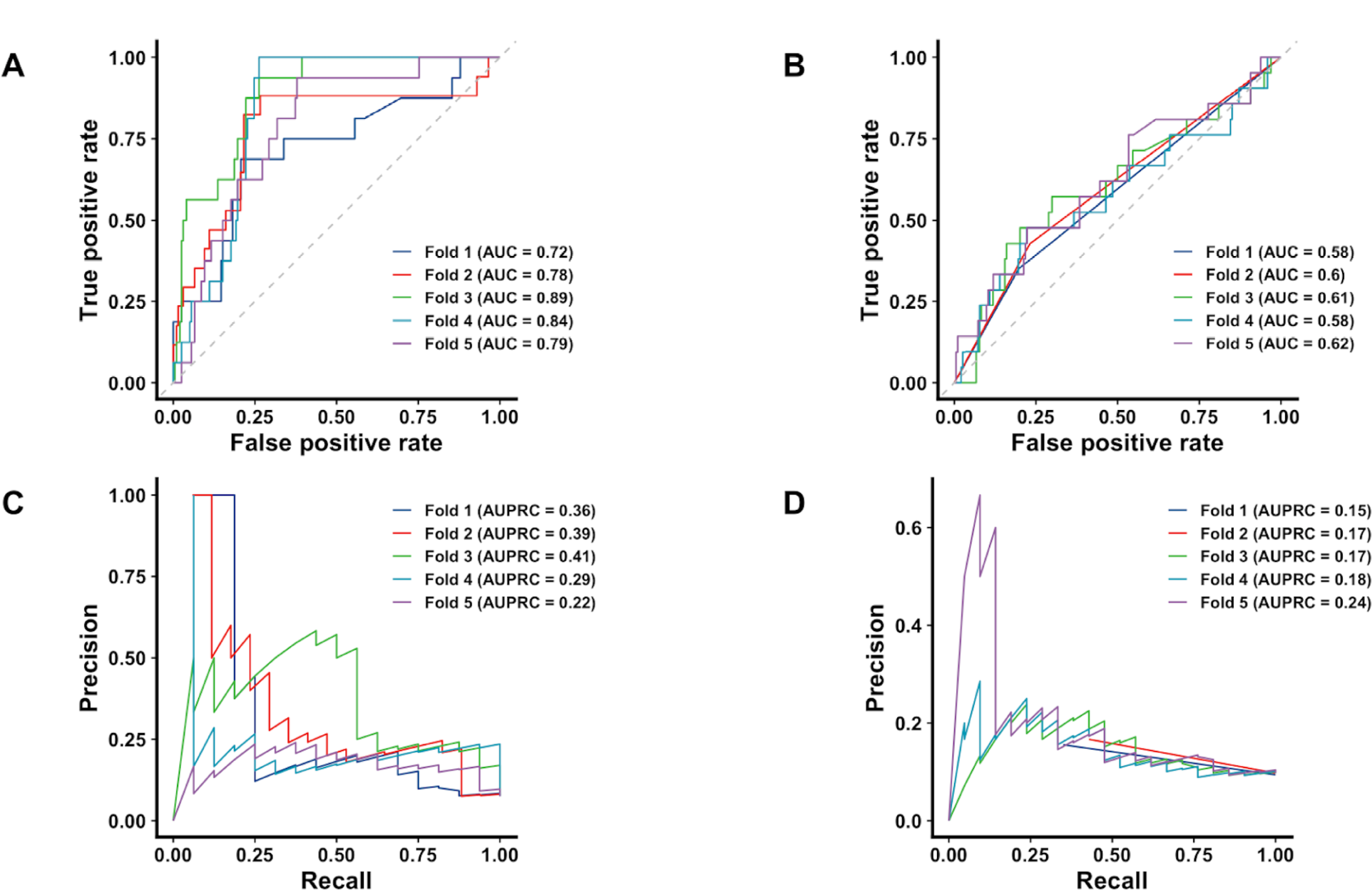
ASD and ID model performance measured by: ROC and PR curves for predicting. (A,C) ASD-related and (B,D) ID-related RBPs. The AUC and AUPRC across different folds are shown as labels.

### Top ASD and ID candidate gene predictions

We examined the top 10 predicted candidate genes for the ASD and ID models (Figure 1C, Table S3). Seven out of the top 10 ASD candidate genes were already present in the SFARI list: *CELF4* (ranked 1^st^), *STXBP1* (ranked 2^nd^)*, ELAVL2* (ranked 3^rd^)*, DYNC1H1* (ranked 5^th^)*, TNRC6B* (ranked 7^th^)*, CHD3* (ranked 8^th^) and *PTBP2* (ranked 9^th^). Importantly, our model predicts three additional ASD genes that are not on this list, but have also been implicated in important brain functions and other neuropsychiatric disorders:

● *CPEB3* (ranked 4^th^) has been shown to be a critical regulator of alternative splicing during neural development and function (Qu et al., 2020), acts as a negative translational regulator of synaptic proteins (Si et al., 2010), and triggers the formation of new synaptic spines (Pavlopoulos et al., 2011). *CPEB3* conditional knockout mice show impaired maintenance of both hippocampal long-term potentiation and hippocampus-dependent spatial memory (Fioriti et al., 2015).
● DDX6 (ranked 6^th^), an RNA helicase, is necessary for neuronal differentiation (Nicklas et al., 2015) and is an ID gene according to Genomics England. *De novo* missense variants have been found in DDX6 in probands with ID (Balak et al., 2019).
● SRPK2 (ranked 10^th^) mediates pre-mRNA splicing, and has been shown to inhibit axonal elongation (Hong et al., 2012). SRPK2 aggravates AD pathogenesis (Wang et al., 2017) and plays an essential role in maintaining the homeostatic balance of synaptic coupling of neuronal networks (Müller et al., 2022).

Out of the top 10 ID candidate genes, *CELF2* (ranked 2^nd^)*, FAM120C* (ranked 3^rd^), and *DDX3X* (ranked 5^th^ and mentioned above in Introduction) are associated with ID according to Genomics England. Our newly identified ID genes have all been found to play important roles in the nervous system and/or to be associated with other neuropsychiatric disorders:

● TNRC6C (ranked 1^st^) is a microRNA that is listed as a SFARI ASD gene pertaining to the respiratory issues faced by the ASD children (Rhine et al., 2022).
● SAFB2 (ranked 6^th^) is highly expressed in the central nervous system and is involved in many aspects of RNA processing, which is crucial for the considerable neuronal cell division that occurs during brain development and the proper function of post-mitotic neurons in the mature brain (Norman et al., 2016).
● RMB17 (ranked 8^th^) is an RBP that represses cryptic splicing of genes. Loss of *Rbm17* in murine Purkinje neurons led to ataxia and rapid degeneration (Tan et al., 2016).
● YTHDC2 (ranked 9^th^) is a “reader” of mRNA N6-methyladenine (m^6^A) modifications (Sokpor et al., 2021) and is associated with ASD according to SFARI.
● FNDC3A (ranked 10^th^) is an RBP located in the Golgi apparatus and has not been extensively studied. However, single-nucleotide polymorphisms in FNDC3A have been associated with ASD (Ro et al., 2013).

Thus, our machine learning model can accurately predict ASD and ID-associated RBPs. In addition, the model can be used to prioritize RBPs that are likely to be important in neuronal function and may play a role in neurodevelopmental diseases.

## DISCUSSION

Here we developed BrainRBPedia, a unique database of RBPs with neurodevelopment-relevant disease and functional annotations. Using the annotations contained in BrainRBPedia, we have further developed a machine learning model to predict and prioritize RBPs that are likely to be associated with ASD and ID. Importantly, half of our top genes predicted for ASD and ID have previously been implicated in these disorders, verifying our model, while half are new and, thus, open up areas of future research into ASD and ID.

Our results showed that RBPs with a very high degree of loss-of-function intolerance and that are upregulated in neuronal development were consistently selected in all cross-validation runs for models of both ASD and ID. Specifically, Extreme pLI, High pLI, and Neuron developmental enrichment (late) were selected for ASD. Despite having more genes annotated as associated with ID than for ASD, our machine learning approaches were considerably less accurate at predicting ID-related RBP genes than for ASD genes. Plausible explanations include that there might be more biological similarity between ASD genes than for ID genes, or that ID annotations are less accurate than ASD annotations.

Overall, RBPs that are highly intolerant to loss-of-function (LoF) protein truncating variations (nonsense, splice acceptor and splice donor variation) are most likely to be associated with ASD and ID. This result is consistent with previous findings that functional mutations in ASD and ID cases are more enriched for loss-of-function intolerant genes, as demonstrated by the RVIS (Residual Variation Intolerance Score) (Samocha et al., 2014). In other psychiatric disorders such as schizophrenia, LoF-intolerant genes are significantly enriched for common risk alleles and account for 30% of the SNP-based schizophrenia heritability in the study (Pardiñas et al., 2018). Furthermore, the effect of RBP dysregulation on target site variants is significantly larger in LoF-intolerant genes compared to the remaining set of genes (Park et al., 2021). In addition, there have been multiple studies showing a link between LoF variants and ASD/ID etiology, many being RBPs. For example, eleven *de novo* loss-of-function mutations in the chromatin remodelling factor CHD8 have been identified in unrelated individuals with ASD (Cotney et al., 2015; Iakoucheva et al., 2019). Cotney et al. showed that the mRNA targets of the CHD8 RBP in the human and mouse developing brains were strongly enriched in ASD risk genes, which also became dysregulated following *CHD8* knockdown. SRRM2, which ranked 25^th^ in our ID predictions, is an RBP involved in mRNA splicing and belongs to the 0.1% most loss-of-function intolerant human protein-coding genes with a pLI score of 1 (Petrovski et al., 2013). From 1,000 probands (i.e., an individual who is affected by or at risk for a genetic condition), Cuinat et al. (Cuinat et al., 2022) found that 22 patients with LoF variants in *SRRM2* display ASD clinical phenotypes including developmental and speech delay. The mutation-phenotype relationships indicate that mutations disrupting specific RBPs may help explain the wide range of symptoms and severity observed in ASD and ID individuals carrying rare mutations.

Our results also indicate that RBPs upregulated in neuron development are also closely linked to ASD and ID. Day 3 (early) and day 7 (late) correspond to the onset of neurite formation and neuronal polarization during neurogenesis, respectively. During neuronal development, neurons must undergo extensive morphological changes after proliferation and migration to acquire complex neuronal processes (axons and dendrites) for synaptic connections and functional neuronal networks. RBPs are essential players in active protein synthesis and maintaining protein homeostasis in the development of neuronal processes (Thelen & Kye, 2020). For example, *RBFOX1* plays a critical role in regulating the alternative splicing of genes critical for neuronal development (Gehman et al., 2011) (Lee et al., 2009) (C. Zhang et al., 2008). Mutations disrupting *RBFOX1* have been associated with ASD, ID, epilepsy, and other neuropsychiatric disorders. *RBFOX1* has a Neuron developmental enrichment (late) value of 2.40, which means it is 5.28 times more enriched at day 7 compared to day 1. As a second example, the RBP CPEB1 co-localizes with beta-catenin mRNA to growth cones of hippocampal neurons and phosphorylation of CPEB1 is correlated with a rapid and exclusive increase in beta-catenin protein accumulation in the growth cones following neurotrophin-3 (NT3) stimulation (Kundel et al., 2009). Further, CPEB1 dysfunction blocks the increase in beta-catenin and causes prolonged exposure to NT3, altering the growth and branching of hippocampal processes (Ivshina et al., 2014).

The main limitation of our model’s predictability comes from the small sample size and the quality of disease annotations in BrainRBPedia. Traditionally, RBPs (canonical RBPs) are defined as proteins that bind to double or single-stranded RNA through RNA binding domains (RBDs) such as RNA recognition motifs (RRMs) and zinc fingers (Moore et al., 2018). Recent advances in proteome-wide approaches for protein-RNA interaction studies have identified many proteins possessing non-canonical RBDs yet endowed with RNA-binding activity (Castello et al., 2012). Here, we have defined RBPs based on the union of two gene lists: genes with canonical RNA binding domains (N = 476) and genes identified experimentally as RBP candidates via interactome capture in HeLa cells (N = 845) (Sundararaman et al., 2016). The small total sample size of 1072 RBPs and the smaller number of disease annotations creates a class imbalance in the dependent variables (i.e. ASD and ID susceptibility). 81 (7.6%) and 104 (9.7%) of the 1072 genes are annotated as associated with ASD and ID, respectively. This means that the model has adequate information about the RBPs that are not associated with a neurodevelopmental disorder (the majority class) but insufficient information about those who are (the minority class), causing the algorithm to be more biased toward predicting the majority class while failing to learn the patterns presented in the minority class. Using a class-specific weighting strategy can help mitigate this limitation; however, including class weights might lead to a strong and systematic overestimation of the probability for the minority class (van den Goorbergh et al., 2022).

In summary, BrainRBPedia provides a unique database useful for downstream efforts to experimentally characterize the roles of RBPs in the etiology of neurodevelopmental disorders. Our machine learning model prioritizes a list of ASD/ID candidate genes and demonstrates that high intolerance to loss-of-function mutations and neuron developmental enrichment are important factors in predicting an RBP’s association to ASD and ID. The prioritized neurodevelopmental disorder candidate genes can now be readily used as experimental targets in follow-up validation studies.

## Supplemental Material

**Table S1.**
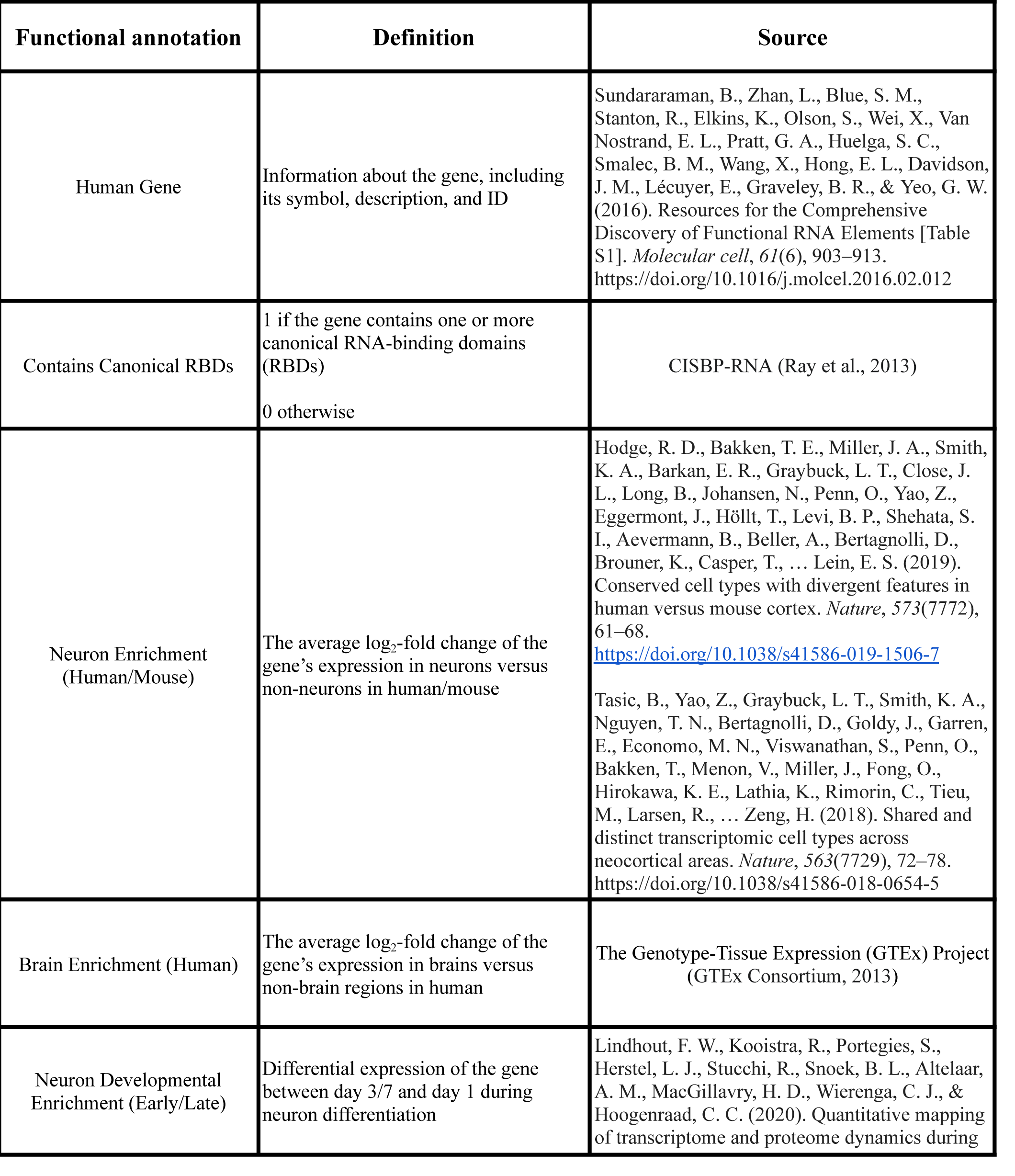
Column Definitions.

**Table S2.**
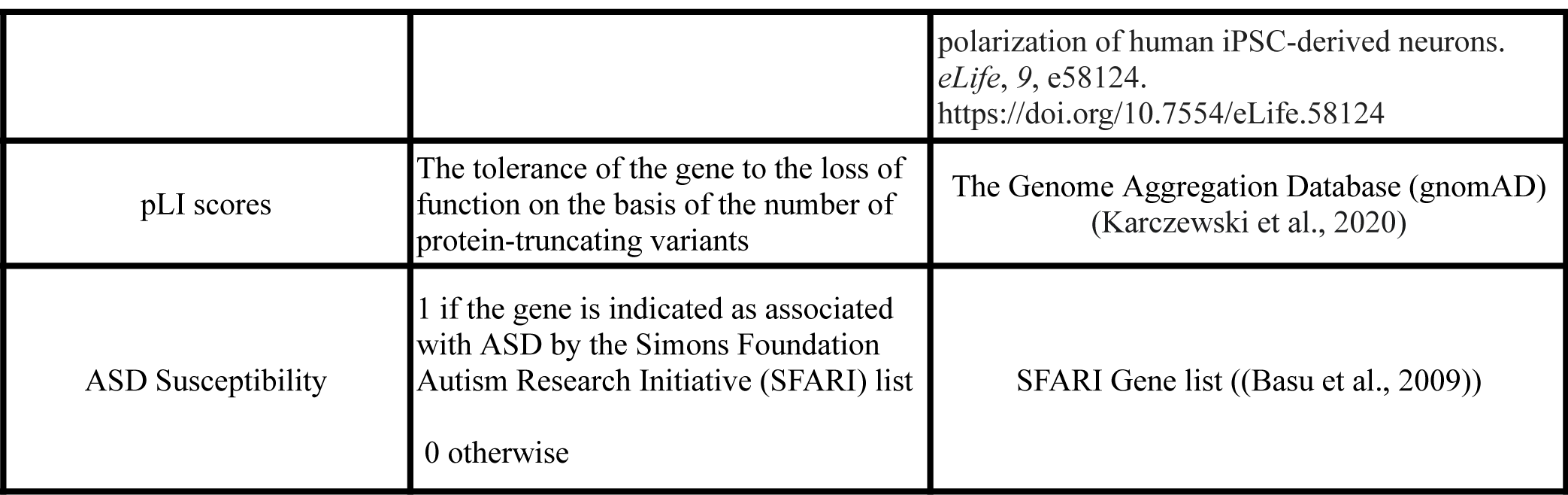
The BrainRBPedia database (see attached)

**Table S3.** Top 10 machine learning model predictions.

| **Ensembl ID** | **HGNC Symbol** | **ASD**<br>**Susceptibility** | **Predicted**<br>**Probability** | **Percentile** |
| --- | --- | --- | --- | --- |
| ENSG00000101489 | CELF4 | 1 | 0.8486 | 100.00 |
| ENSG00000136854 | STXBP1 | 1 | 0.8185 | 99.91 |
| ENSG00000107105 | ELAVL2 | 1 | 0.8114 | 99.81 |
| ENSG00000107864 | CPEB3 | 0 | 0.8012 | 99.72 |
| ENSG00000197102 | DYNC1H1 | 1 | 0.7970 | 99.62 |
| ENSG00000110367 | DDX6 | 0 | 0.7664 | 99.53 |
| ENSG00000100354 | TNRCD6B | 1 | 0.7603 | 99.44 |
| ENSG00000170004 | CHD3 | 1 | 0.7592 | 99.35 |
| ENSG000000117569 | PTBP2 | 1 | 0.7569 | 99.25 |
| ENSG000000135250 | SRPK2 | 0 | 0.7487 | 99.16 |

| **Ensembl ID** | **HGNC Symbol** | **ID Susceptibility** | **Predicted Probability** | **Percentile** |
| --- | --- | --- | --- | --- |
| ENSG00000078687 | TNRC6C | 0 | 0.6140 | 100.00 |
| ENSG00000048740 | CELF2 | 1 | 0.6089 | 99.91 |
| ENSG00000184083 | FAM120C | 1 | 0.6070 | 99.81 |
| ENSG00000075856 | SART3 | 0 | 0.6057 | 99.72 |
| ENSG00000215301 | DDX3X | 1 | 0.6045 | 99.63 |
| ENSG00000130254 | SAFB2 | 0 | 0.6044 | 99.53 |
| ENSG00000134744 | ZCCHC11 | 0 | 0.6031 | 99.44 |
| ENSG00000134453 | RBM17 | 0 | 0.6024 | 99.35 |
| ENSG000000047188 | YTHDC2 | 0 | 0.6018 | 99.25 |
| ENSG000000102531 | FNDC3A | 0 | 0.6013 | 99.1 |

**Table S4.**
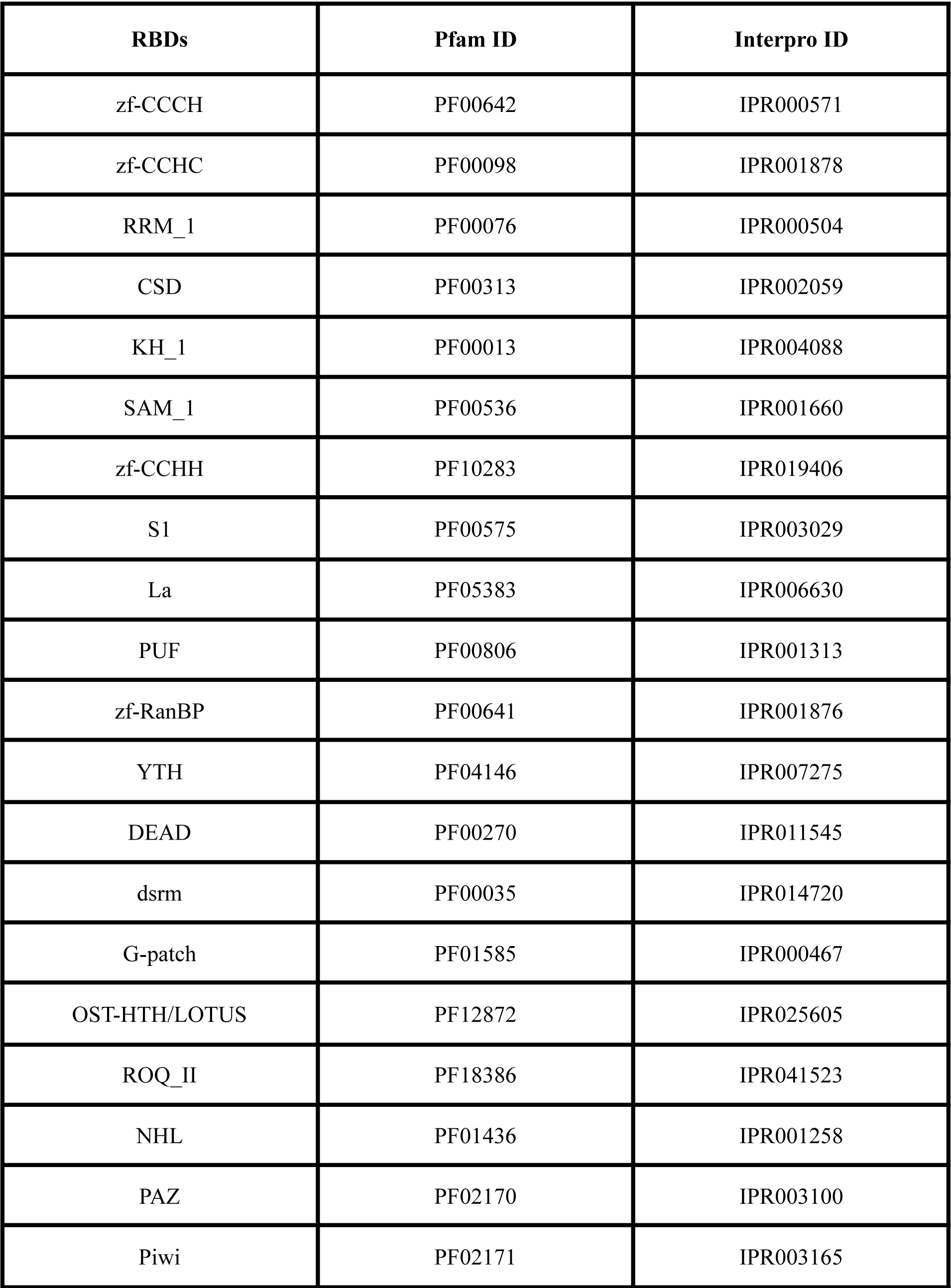
List of canonical RNA-binding domains.

**Table S5.** Odds ratios from ASD and ID machine learning models (see attached)

